# ISRES+: An improved evolutionary strategy for function minimization to estimate the free parameters of Systems Biology models

**DOI:** 10.1101/2023.04.29.538818

**Authors:** Prasad U. Bandodkar, Razeen R. Shaikh, Gregory T. Reeves

## Abstract

Model development is essential to gain a mathematical understanding of the underlying phenomena in systems biology. In most models, it is typically hard to estimate the values of the biophysical/phenomenological parameters that characterize the model. The parameters are estimated by minimizing a function that reduces a measure of the error between model predictions and experimental data. In this work, we build on an algorithm for function minimization proposed by Runnarson and Yao, named Improved Evolutionary Strategy by Stochastic Ranking (ISRES), that finds a best-fit individual by evolving a population in the direction of minimizing error by using information at most from a pair of individuals in any generation to create a new population. Our algorithm, named ISRES+, builds on it by combining information from all individuals across the population and across all generations to gain a better sense of direction to evolve the population. ISRES+ makes use of the additional information generated by the creation of a large population in the evolutionary methods to approximate the local neighborhood around the best-fit individual using linear least squares fit in one and two dimensions. We compared the performance of the two algorithms on three systems biology models with varying complexities and found that not only does the ISRES+ lead to fitter individuals, but it also leads to a tighter distribution of fittest individuals over successive runs.

## Introduction

Here we report an improved algorithm for the optimization of continuous variable non-linear functions. Typically, in systems biology, information from large volumes of data from sequencing to spatial gene expression is condensed to build mathematical models that can describe the data concisely as well as make predictions to guide further examination. These models tend to have several parameters that may be phenomenological or mechanistic. Even in the mechanistic models, it is usually difficult to measure all the biophysical parameters that characterize the model to similar precision. Such problems can be represented as a function optimization problem where the objective is to estimate the values of the parameters that reduce a measure of error between model predictions and experimental data. These models are usually expressed in the form of coupled ordinary or partial differential equations, where the state variables are protein/mRNA concentrations which are then compared to experimentally obtained data.

In this paper, we report an algorithm that builds on the Improved Evolutionary Strategy by Stochastic Ranking (ISRES) (Runarsson & Yao, 2005). If a vector of model parameters can be thought of as an individual in a population, the algorithm is basically a strategy for the evolution of a population of randomly chosen individuals in search space to a population that is tightly clustered around the fittest individual. The fittest individual at the end of the evolution would represent the final vector of model parameters that best fits the model predictions to the data. In a constrained optimization problem, the optimum search direction is determined by using an objective function that measures the error between model predictions and data and a penalty function that measures the feasibility of the solution. The authors use stochastic ranking to create a balance between the objective and penalty functions to rank the population in a constrained optimization problem. In every generation, a fixed number of highest ranked individuals are selected to produce offspring to create a new population. This is done partially by recombination, where the fittest individual mates with the other selected individuals, and partially by mutation of the constituent parameters of the selected individuals. Thus, while the search direction is biased to probe the region around the fittest individual in every generation, the search space around all selected individuals is also explored. In this manner, the algorithm converges to a region around a function minimum and continues to improve its estimate by exploring successively smaller regions of the search space by creating tightly distributed population clusters.

Our method takes advantage of the excess information generated by the Evolutionary Strategy (ES). In any generation, several trials are held to explore the fitness landscape but only a few are selected to create progeny for the next generation. The idea is to use information only from the population but also from the lineage of all individuals in the population to better understand the fitness landscape. We use linear least squares fit to fit either a linear or a quadratic model to the data in every generation. The linear least squares fit coefficients can then be correlated to the coefficients of a Taylor series expansion to construct approximate values of the gradient and the Hessian. A linear model is used to build an estimate of the gradient, which is used to perform an approximate gradient descent step and a quadratic function model is used to build an estimate of the Hessian which is used to perform an approximate Newton’s method of function optimization step. In every generation, in addition to the recombination and mutation contributions, new individuals are created from top ranked individuals by the linear least squared fits and added to the population. Note that these methods add only minimal overhead to the creation of new offspring, since the neither the gradient nor the Hessian is really computed. Both are estimated using the coefficients of the linear least squares fit. In order to maintain maximum progress, the fitting is done in the local neighborhood of the fittest individual found in all generations. The population size is kept constant in every generation, by adding these individuals at the expense of some of the mutation contributions of the lowest ranked individuals. The objective of these methods is to provide a better estimate of the search direction and complements the recombination method of the evolutionary strategy (ES).

We applied our algorithm to three systems biology models and found that when the linear least squares fit based methods contribute ∼2% to a new population in every generation, there was a higher probability of converging to a better solution for the same set of hyperparameters that characterize the ES, such as the population size and the number of generations. In addition, the final distribution of best parameter values was narrower with the addition of the linear least squares fit contributions to ISRES as compared to without it.

## Methodology

In this study we investigate three systems biology models described by ODEs and compare the performance of the ISRES+ algorithm against the original ISRES (Runarsson & Yao, 2005). The models vary in their complexity, in terms of the number of parameters, the number of state variables and the number of spatiotemporal points (Table 1). The objective function is defined as the sum of squares error according to the following equation,

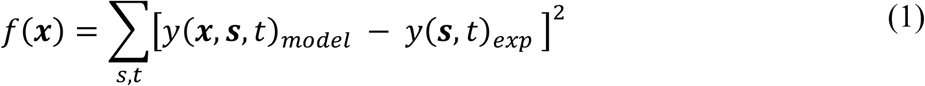

where ***x*** is the parameter vector, ***s*** and *t* are the spatial and time coordinates. When applicable, constraints on the feasible regions are expressed as,

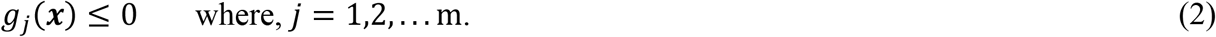

where *m* is the number of constraints. The fitness function is then represented as,

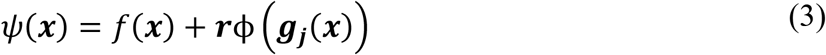

where the penalty function *ϕ*(***x***) is a quadratic penalty function defined as

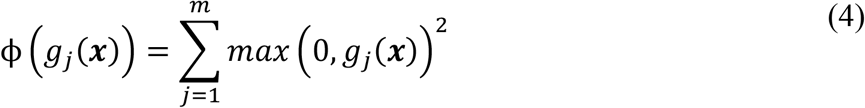

and ***r*** is a vector of penalty coefficients that regulates the dominance between the objective and the penalty functions in determining fitness. ISRES uses stochastic ranking to handle constraints in a way that it neither under-penalizes (low ***r***) nor over-penalizes (high ***r***) the fitness function. The primary aim of stochastic ranking is to maintain a balance between the objective and penalty functions ensuring that an optimum search direction is maintained not just in the overall search space but in the feasible search space (Runarsson & Yao, 2000).

**Table 1.** Model complexity across the three systems biology models investigated in this study.

| Model | # free parameters | # optimization points | # system variables |
| --- | --- | --- | --- |
| Model 1: Dl/Cact model | 15 | ~12000 | ~300 |
| Model 2: Smad signaling | 10 | ~50 | 25 |
| Model 3: Gap gene circuit model | 44 | ~1800 | 228 |

The search boundaries for the parameters of each of the models ranges from 1e-4 to 1e+4, unless it is known from experimental observations or scaling arguments that a parameter is restricted to vary in a different range. The objective is to find a value for the parameter vector ***x***, that the algorithm would characterize as the fittest individual found over the course of evolution across all generations.

## Algorithm

ISRES+ builds on the ISRES algorithm, proposed by Runnarson & Yao, for minimization of continuous variable non-linear functions (Runarsson & Yao, 2005). Fig. 1 describes the ISRES+ algorithm that includes steps from ISRES. In the original ISRES algorithm, a population of *λ* individuals is initialized from an *n*-dimensional uniform distribution, where an individual is considered as a set of *n* parameters that characterizes the model. The population is then ranked using stochastic ranking by maintaining a balance between the objective and penalty functions in a way that it only slightly favors the feasible search space (which is defined as the region of search space in which the penalties are satisfied). If it is an unconstrained optimization problem, then only the objective function is used for ranking. In every generation, *μ*(< *λ*) top-ranked individuals are selected for evolution to the next generation while the rest are discarded. The selected individuals produce offspring to populate the next generation by two methods. In the first method, the top-ranked individual mates with the other parents to produce *μ* − 1 offspring by the process of recombination according to the equation,

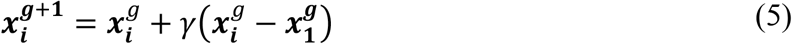

where, *i* = (2, …, *μ*), *g* refers to the generation, and *γ* is the recombination parameter. In the second method, each of the selected parents are repeated rank-wise to fill the remaining *λ* − *μ* + 1 population and the constituent parameters of an individual are subject to random mutations according to a gaussian probability distribution, according to the equation,

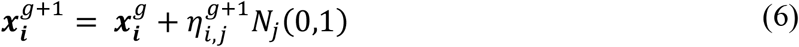

where, *i* = (*μ* + 1, …, *λ*), *j* = (1, …, *n*), *N*_*j*_ is a normally distributed random number for each parameter, and

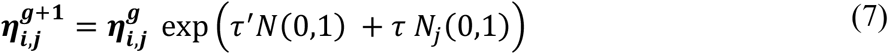

where, ***η***_***i***,***j***_ is a strategy parameter which is essentially a measure of mean step sizes for each model parameter to take to maintain maximum progress to reach the function minimum, 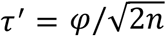 is the learning rate for an individual, 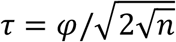 is the learning rate for each parameter, and *N*(0,1) is a normally distributed random variable vector (Beyer, 1995; Schwefel & Rudolph, 1995). The variable *φ* is the expected rate of convergence and is usually set equal to 1. The mutation strategy is self-adaptative, as the strength of mutation is independent of the rest of the population and depends only on its parents’ strategy parameters, and is non-isotropic, as each parameter of each individual is multiplied by a distinct random number. At the end of each generation, mutation strength is reduced by exponential smoothing according to equation,

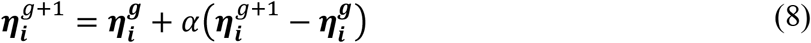

where, *i* = (1 … *λ*) and *α* is the smoothing factor (Runarsson, 2002). The ratio of recombination contributions to that of mutation is kept at ∼1/7 to roughly ensure an equal probability of success in finding a fit individual by the two methods (Runarsson & Yao, 2005).

**Figure 1.**
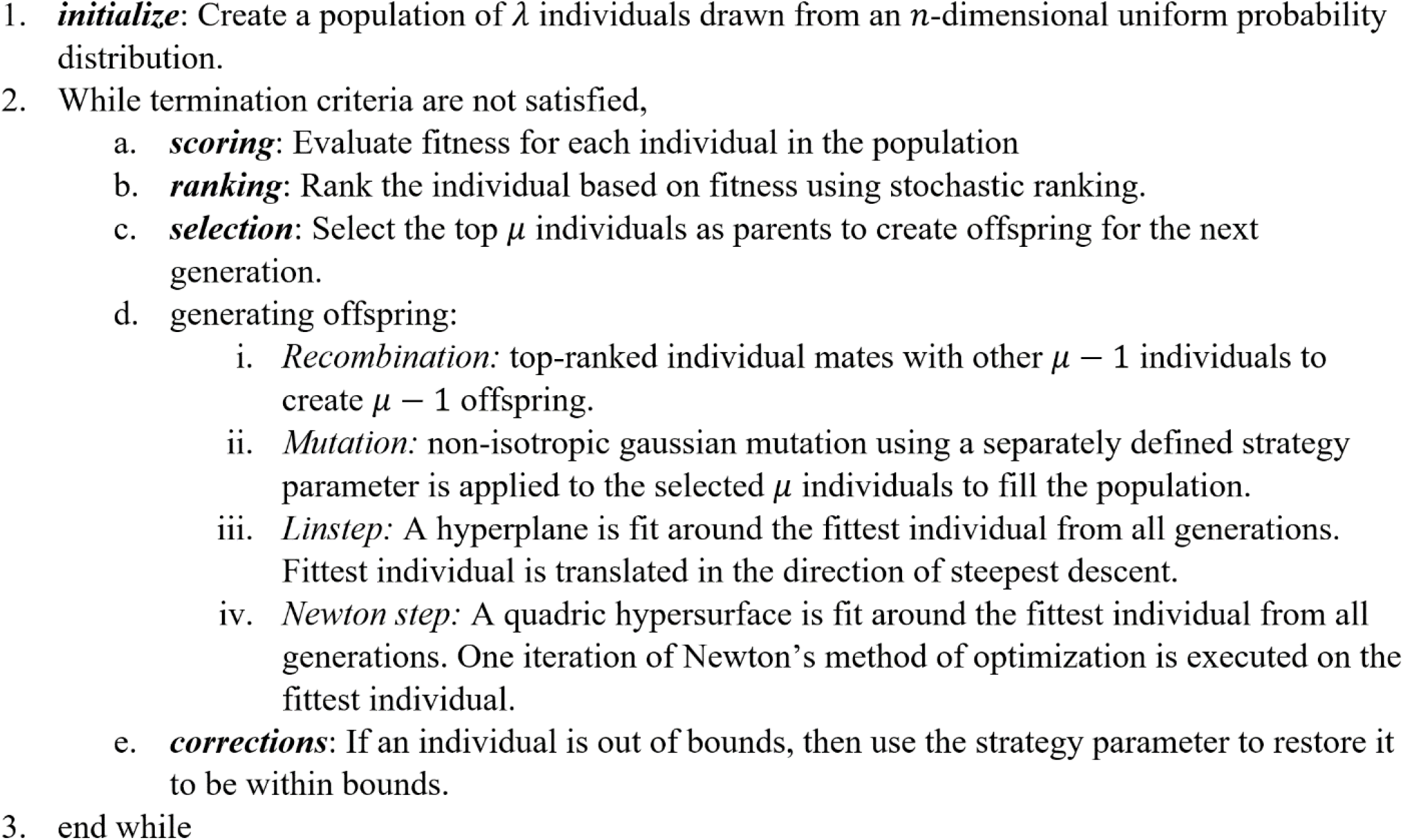
The ISRES+ algorithm

ISRES+ extends the original ISRES algorithm by obtaining directions to evolve the individuals in by fitting the fitness landscape in the local neighborhood of the best individual to either first or second order polynomials of the parameters. To fit a first order polynomial (hyperplane), a minimum of *n* + 1 feasible individuals are required. First, all feasible individuals are sorted by Euclidean distance from the fittest one. The fittest individual along with the cluster of the closest *n* individuals around it are then used to fit a hyperplane, which is then used to determine the gradient **v** = *df*/*d****x***for all individuals in the cluster, where *f* is the fitness and ***x***is the parameter vector. The obtained gradient approximates the direction of steepest descent.

Individuals from the cluster are then translated along the steepest direction according to the equation,

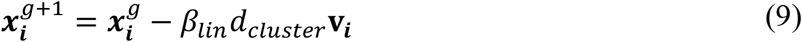

where *i* = (*λ* − *nlin*),.., *λ*; *nlin* is the number of Linstep contributions to the population; *β*_*lin*_ ≥ 1, and *d*_*cluster*_ is the diameter of the cluster bubble of individuals that were used to determine the gradient. This method, henceforth called Linstep, is an approximate steepest descent step.

To fit a second order polynomial, the fittest individual along with a cluster of the closest (*n*^2^ + *n*)/2 individuals are used. The coefficients of the second order polynomial are used to construct an approximate Hessian, **H** = *d*^2^*f*/*d****x***^2^, and a new value of the gradient. These approximations are then used to perform a single iteration of the Newton’s method of function optimization on the individuals from the cluster, according to the equation,

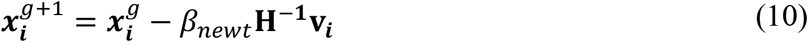

where, *i* = (*λ* − *nlin* − *nnewt*),.., (*λ* − *nnewt*) and 0 < *β*_*newt*_ ≤ 1. In each case, a minimum number of individuals are chosen to determine these directions, and if the resulting matrix of the gradient or the Hessian is ill-conditioned, then more individuals are used. The individuals contributed by Linstep and Newton step are added to the population at the expense of mutations around the least fit parents according to the ES, thus maintaining the size of the population *λ*.

The population continues to evolve by each of the four aforementioned methods for a pre-determined number of generations. The algorithm outputs the fittest individual found in all generations and the value of its objective function, along with some statistics about the evolution.

## Results

We applied both ISRES+ and ISRES to three systems biology models to compare how each algorithm behaves over successive generations and fares against each other. The first model, referred to as the Dl/Cact model, describes a morphogen system model that patterns the dorsalventral axis of early *Drosophila* embryos (Kanodia et al., 2009; O’Connell & Reeves, 2015; Schloop et al., 2020). The second model, referred to as the Smad signaling model, describes TGF-*β* induced Smad2 signaling in HaCat cells (Schmierer et al., 2008). The third model, referred to as the gap gene circuit model, describes the patterning of the anterior-posterior axis of early *Drosophila* embryos (Jaeger, 2009; Manu et al., 2009). For the evolutionary strategy part of the algorithm, we used the recommendations for the settings of the hyperparameters from the original ISRES algorithm proposed by Runnarson & Yao in all cases (Runarsson & Yao, 2005). For every combination of hyperparameters, we ran each of the algorithms at least 50 times. The comparisons in all the rest of the plots were made with the modified version of the original ISRES algorithm.

For the Dl/Cact model, the number of Linstep and Newton step contributions to the population were varied and the performance of ISRES+ was compared against ISRES. In Fig 2, the value of error for the fittest individual across at least 50 independent simulations is plotted in the first two rows of plots while the last row of plots are histograms of the best individual obtained at the end of the simulation. The three plotlines in the first two rows of plots indicate the 25^th^, 50^th^ and 75^th^ percentile of the minimum value of error (inverse fitness) for every generation across the >50 independent simulations. The histogram plots, however, represent all data. In Figs. 2A-C, only Linstep is active during the entire duration of the simulation and contributes two individuals to the population in every generation, while in Fig. 2D-F, only Newton step is active and contributes one individual. In Figs. 2G-I, both Linstep and Newton step are active with the former contributes two individuals while the latter contributes one individual to the population in every generation. As seen in Fig 2A-B, when only Linstep is on, the ISRES+ performs slightly better than ISRES at the 25^th^ and 50^th^ percentile mark. The histogram of the final error values in Fig. 2C indicates that, across the 50 independent runs ISRES+ performs only slightly better than ISRES. As seen in Fig. 2D-F when only Newton step is on, the final performance improves slightly, but the final distribution of individuals forms a tighter cluster. When both Linstep (*n*_*lin*_ = 2) and Newton step (*n*_*newt*_ = 1) contribute to the population, the breadth of the final distribution is intermediate to the cases when only one of the two methods is active but is better than either case. For the Dl/Cact model, Linstep appears to improve the overall performance while Newton step assists in getting a tighter final distribution. Note that the 75^th^ percentile plotline for ISRES+ follows that of ISRES, when only Linstep is active (Figs. 2A-C), and the plotline converges markedly at G∼=300 when only Newton step is active. When both Linstep and Newton step are active, the converging behavior is observed later at G∼=500. For the Dl/Cact model, while Newton step seems to assist in the converging behavior that leads to a tighter final distribution. (Fig. 2F), both methods are required to also obtain a better final distribution (Fig. 2I). We have observed that, regardless of the specific combination of hyperparameters, Linstep and Newton step take the algorithm to better solutions faster and the overall distribution of the final parameter values over several independent runs is much tighter.

**Figure 2.**
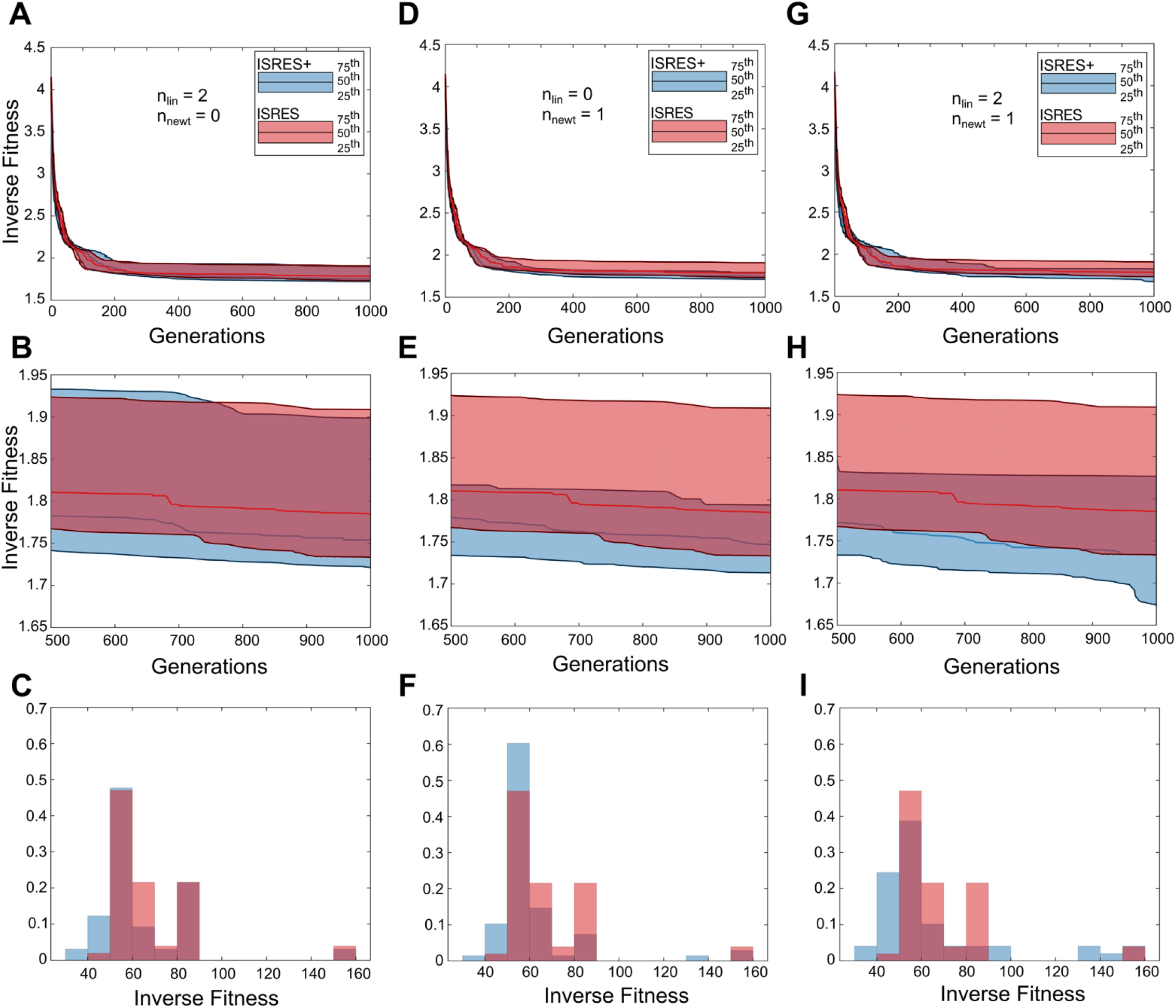
ISRES+ v/s ISRES comparison for the Dl/Cact model.

For the Smad signaling model, we found that the model had at least two local minima, one around *error* ≅ 6 and the other around *error* ≅ 0.5. From Figs. 3A,C, and E, we find that both algorithms converge to either one of the two minima fairly quickly (by *G* < 100). However, fewer than 25% of the ISRES simulations converge to the lower minimum (Figs. 3B,D,F), which is why it appears that ISRES is unable to find the lower minimum in Figs. 3A,C,E, since these plots only represent the data between the 25^th^ and 75^th^ percentile and do not represent the entire >50 simulation runs. In Figs. 3B, D, and F, the histograms of the final distribution of error values normalized by probability are plotted. It can be seen that while ISRES does find the lower value of minimum error, it does so less often than ISRES+ for the various combination of hyperparameters. In particular, when *β*_*lin*_ > 1, regardless of the specific values of the other hyperparameters, ISRES+ not just outperformed ISRES but was able to find a better minimum more often (Fig. 3). Also, ISRES+ performs better when *β*_*lin*_ = 2 than when *β*_*lin*_ = 1.5, indicating that the bulk of the gains obtained may be attributed to Linstep. Thus, for the Smad signaling model, higher values of *β*_*lin*_ favor better final distributions over several independent runs.

**Figure 3.**
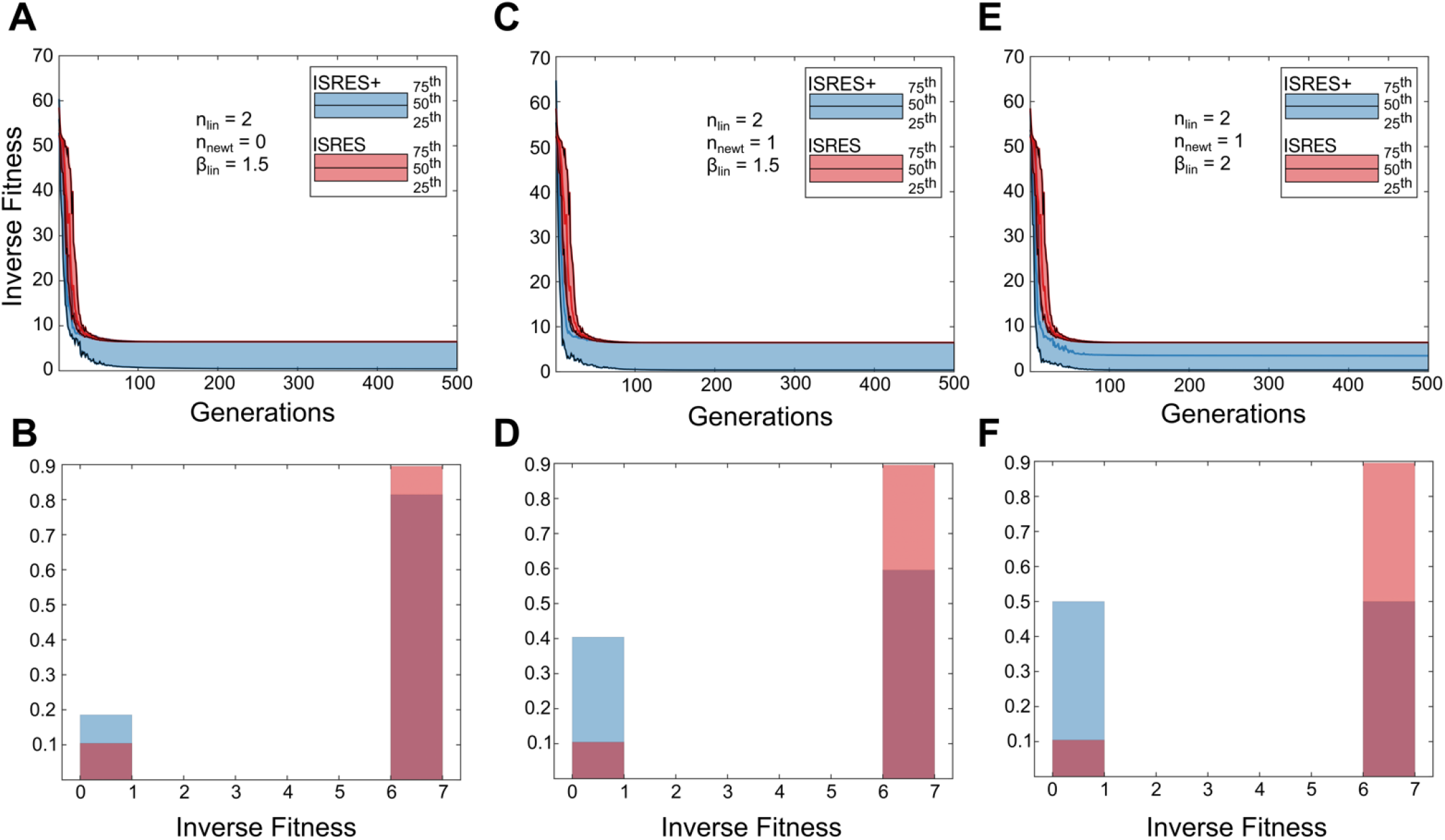
ISRES+ v/s ISRES comparison for the Smad signaling model

For the gene circuit model, we varied the generation at which each of Linstep and Newton step is switched on or off (Fig. 3). Note that in all configurations of hyperparameters, *n*_*lin*_ = 2 and *n*_*newt*_ = 1. In Fig.4A,B, Linstep is active from *G* = 1 − 1000, while Newton step is active from *G* = 1001 − 3000. Since the model is relatively more complex, the idea was to be able to provide the evolutionary part of the algorithm enough power in the search. Linstep contributions are expected to dominate earlier in the search strategy, when the population is more randomly distributed, while Newton step is expected to perform better later since the population may be expected to be closer to a function minimum. For the configuration in Figs. 4A,B, we find only a small improvement over ISRES. When *β*_*lin*_ = 2, the same configuration performs much better as shown in Figs. 4C,D. In Figs. 4E,F, when *β*_*lin*_ = 2 and when both Linstep and Newton step are active during the entire duration of evolution, ISRES+ performs significantly better than ISRES.

**Figure 4.**
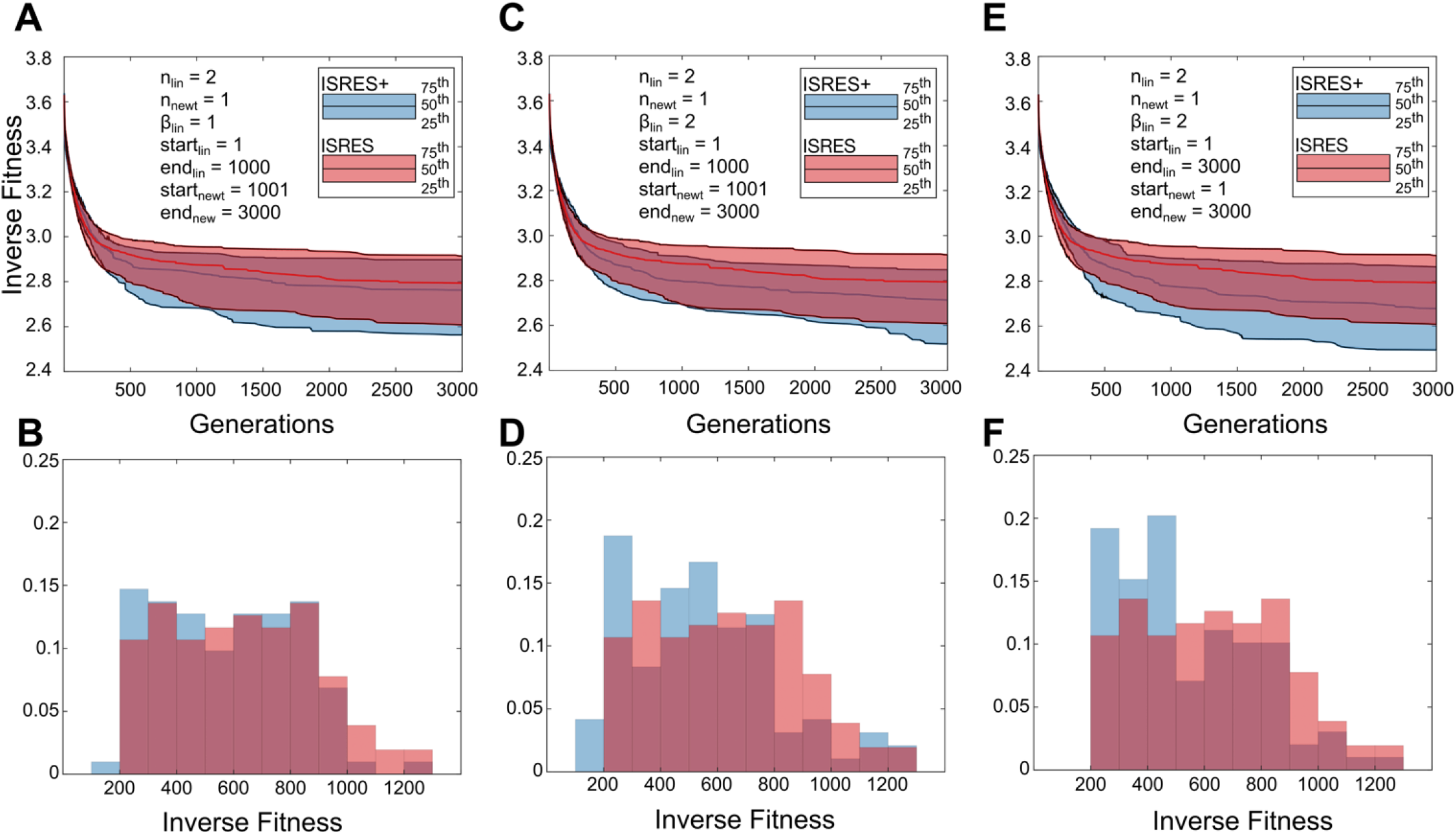
ISRES+ v/s ISRES comparisons for the gap gene circuit model.

## Discussion & Conclusions

The performance of a function optimization algorithm rests on its ability to handle constraints while simultaneously minimizing the objective function. In ISRES, a balance is maintained between the objective and penalty function values by stochastically ranking the individuals in a way that it only slightly favors the feasible search space, as demonstrated here by the Dl/Cact model. Ranking is performed by using a bubble-sort-like procedure, where the values of the penalty functions of adjacent individuals are compared and their ranking is swapped if the lower ranked individual has a lower penalty. A swapping could also occur, if the objective function value of the lower ranked individual is lower to a certain probability (0.45). Thus, stochastic ranking attempts to balance the dominance of either function in determining the fitness, and consequently ranking. Note that the objective is to be able to maintain maximum progress not just toward a function minimum, but to a feasible function minimum. We found that, in the Dl/Cact model, while in earlier generations, the population has a near equal mix of feasible and non-feasible individuals, the number of feasible individuals in the population steadily increases as the algorithm moves towards a region of parameter space that is both fit and feasible. Progress in function minimization is also influenced by the search methods, namely, recombination and mutation. In recombination, the top-ranked individual of a particular generation mates with the other parents to create offspring for the next generation. In this method, the search space between the line connecting the top-ranked individual and the other parents is explored. In other words, information is shared between a pair of individuals and the overall search space is moved toward the fitter one. In mutation, a self-adaptive method is used where the mutation strength is defined by a strategy parameter that is inherited from the parent multiplied with a random number drawn from a Gaussian probability distribution. The strategy parameter is steadily decreased over generations in a way that depends only on the lineage, and not on the rest of the population. Thus, neither the mutation strategy parameter nor its steady decrease over the course of evolution is dependent on the population. The rationale behind the approach is to avoid considering very different individuals from the population while evaluating mutation strength in any generation. Therefore, in the mutation method of creating progeny, no information is shared between individuals in the population.

Since the entire population in an evolutionary algorithm steadily moves from a region spanning the entire the search space to a relatively small region around a function minimum, there is an abundance of information available about the fitness landscape not only across the population but also across generations. Our method seeks to take advantage of this information in understanding the features of the fitness landscape, to derive better search directions to evolve the population. The fitness landscape is fit in the local neighborhood of the best individual to either first or second order polynomials of the parameters using linear least squares fitting.

In the first order fit, used by Linstep, the closest *n* individuals to the fittest individual in all generations is used to fit a hyperplane. The polynomial coefficients of the hyperplane are then used as an approximation of the steepest descent direction which would be analogous to a first order Taylor series expansion. In Linstep, this direction is used to translate a small number of the fittest individuals (1-3), in the direction of the steepest descent estimate, creating new offspring. Linstep is expected to work better in the earlier generations when the best parameter set may be far from a minimum, in which case only a general direction of descent is needed, rather than a Newton-type step, which would afford a more precise move towards a minimum. On the other hand, in later generations, when the population may be closer to a minimum, Linstep could potentially overshoot a minimum basin in a phenomenon known as gradient hemistitching. In fact, Linstep is likely more prone to this issue than traditional gradient descent methods, as the parameter sets that are used to generate the hyperplane approximation may lie on either side of a minimum basin, which would result in a highly inaccurate gradient approximation.

In the second order polynomial fit, used by Newton step, the closest (*n*^2^ + *n*)/2 individuals along with the fittest one, are used to fit a quadric hypersurface paraboloid in *n* dimensions. The polynomial coefficients of the paraboloid are then used to construct an approximate Hessian and a new value of the gradient for each individual, which would be analogous to a second order Taylor series expansion. In Newton step, the estimate of the Hessian and the gradient are used to perform a single iteration of Newton’s method of optimization on a handful of individuals (1-3) that were used to make the estimate. It is interesting to note that performing a full Newton’s step (*β*_*newt*_ = 1 in eqn. 10) takes all individuals considered in the Hessian calculation to the same point in search space. Therefore, this method could either contribute a single individual from a full Newton step or several with varying step sizes. Newton’s step is expected to work better in later generations as the population converges on a function minimum in the fitness landscape.

In all three Systems Biology models, we find that small contributions from Linstep and Newton step to the population biases the search in a way that it converges to a more fit region of parameter space faster and more often. Also, the final distribution of individuals obtained across multiple independent runs is much tighter as compared to ISRES. The majority of the initial gains may be attributed to Linstep, while Newton step seems to be responsible for ensuring that the algorithm converges to the same region in function space over multiple runs. It is also observed that regardless of the specific values of the hyperparameters, small contributions of the two methods to the population in every generation leads to better overall performance. This may be attributed to the fact that Linstep and Newton step attempt to direct the strategy to search around the fittest individual found so far, even if the ES methods have shifted focus to exploring other regions that might neither have lower penalty values nor lower objective function values. Since the methods focus on the landscape local to the fittest individual in all generations, there is mixing of information not only from within the population but also between generations. In some sense, the entire evolutionary history is used to develop a better understanding of the functional landscape around the fittest individual found so far and optimal search directions are obtained in a deterministic manner.

