## Supplemental Information for "ISRES+: An improved evolutionary strategy for function minimization to estimate the free parameters of Systems Biology models"

### 1. Model Description

#### 1.1 Dl/Cact model

##### 1.1.1 Description

The Dl/Cact model describes the formation of the Dorsal (Dl) gradient that patterns the dorsal-ventral axis in early *Drosophila* embryos (Asafen et al., 2020; Carrell et al., 2017; Kanodia et al., 2012; O'Connell & Reeves, 2015). Dorsal is a maternal morphogen that is initially uniformly expressed in the early embryo. Dl binds to another maternally deposited protein, Cactus (Cact), in the cytoplasm that inactivates it. The Dl/Cact complex binds to Toll receptors resulting in its dissociation leading to free Dl. Since Toll is distributed asymmetrically along the dv axis, the Dl gradient is formed with high concentrations along the dorsal midline. The Dl gradient then activates genes in a concentration dependent manner and patterning the DV axis into three tissue types – mesoderm, neuroectoderm, and dorsal ectoderm.

##### 1.1.2 Mathematical model

The mathematical equations of the model may be represented as follows,

$$\frac{d\left[V_{nuc}C_{d,nuc}^{h}\right]}{dt} = A_{nuc}\left(k_{in,d}C_{d,cyt}^{h} - k_{out,d}C_{d,nuc}^{h}\right) - V_{nuc}\left(k_b C_{d,nuc}^{h} C_{c,nuc}^{h}\right) \tag{1}$$

$$\begin{aligned}\frac{d\left[V_{cyt}C_{d,cyt}^{h}\right]}{dt} = {} & A_{cyt}\Gamma_d\left(C_{d,cyt}^{h-1} - 2C_{d,cyt}^{h} + C_{d,cyt}^{h+1}\right) + V_{cyt}\left(\frac{k_d(x)C_{dc,cyt}^{h}}{\kappa + C_{dc,cyt}^{h}}\right. \\ & \left. - \; k_b C_{d,cyt}^{h} C_{c,cyt}^{h}\right) - A_{nuc}\left(k_{in,d}C_{d,cyt}^{h} - k_{out,d}C_{d,nuc}^{h}\right)\end{aligned} \tag{2}$$

$$\frac{d\left[V_{nuc}C_{dc,nuc}^{h}\right]}{dt} = A_{nuc}\left(k_{in,dc}C_{dc,cyt}^{h} - k_{out,dc}C_{dc,nuc}^{h}\right) + V_{nuc}\left(k_b C_{d,nuc}^{h} C_{c,nuc}^{h}\right) \tag{3}$$

$$\frac{d\left[V_{cyt}C_{dc,cyt}^{h}\right]}{dt} = A_{cyt}\Gamma_{dc}\left(C_{dc,cyt}^{h-1} - 2C_{dc,cyt}^{h} + C_{dc,cyt}^{h+1}\right) - V_{cyt}\left(\frac{k_d(x)C_{dc,cyt}^{h}}{\kappa + C_{dc,cyt}^{h}} - k_b C_{d,cyt}^{h}C_{c,cyt}^{h}\right) - A_{nuc}\left(k_{in,dc}C_{dc,cyt}^{h} - k_{out,dc}C_{dc,nuc}^{h}\right) \tag{4}$$

$$\frac{d\left[V_{nuc}C_{c,nuc}^{h}\right]}{dt} = A_{nuc}\left(k_{in,c}C_{c,cyt}^{h} - k_{out,c}C_{c,nuc}^{h}\right) - V_{nuc}\left(k_b C_{d,nuc}^{h}C_{c,nuc}^{h}\right) \tag{5}$$

$$\frac{d\left[V_{cyt}C_{c,cyt}^{h}\right]}{dt} = A_{cyt}\Gamma_{c}\left(C_{c,cyt}^{h-1} - 2C_{c,cyt}^{h} + C_{c,cyt}^{h+1}\right) + V_{cyt}\left(\frac{k_d(x)C_{dc,cyt}^{h}}{\kappa + C_{dc,cyt}^{h}} - k_b C_{d,cyt}^{h}C_{c,cyt}^{h} - k_{deg}C_{c,cyt}^{h}\right) - A_{nuc}\left(k_{in,c}C_{c,cyt}^{h} - k_{out,c}C_{c,nuc}^{h}\right) + P_c \tag{6}$$

where, subscripts *nuc* and *cyt* represent nucleus and cytoplasm; subscripts *d*, *c*, and *dc* represent the species Dl, Cact and the Dl/Cact complex; A, V, and C represent area, volume, and concentrations; $\Gamma$ represents intercompartmental exchange rates; $k_d(x) = k_d^{max} exp\left({}^{x}/_{\phi}\right)^2$, $k_b$, $k_{deg}$, and κ represent gaussian Toll-mediated rate constant, the Dl/Cact binding constant, the degradation rate constant for Cact, and Michaelis Menten constant for the dissociation of Dl/Cact complex respectively; $k_{in}$ and $k_{out}$ represent nuclear import and export rates respectively; and $P_c$ represents rate of production of Cact.

##### 1.1.3 Simulation Conditions

The Dl/Cact model was run according to O'Connell and Reeves, 2015, where a similar model was first introduced. Live imaging of Dl was used as the experimental data and the corresponding spatio-temporal grid generated was implemented in the model. Note that the number of spatio-temporal points change with nuclear cycles. The model data was rescaled to that of the experimental data to ensure appropriate error calculations.

##### 1.1.4 Model Results

The following figure shows the concentration of Dl ($C_{d,nuc} + C_{d,cyt}$) plotted against simulated data at the ventral ($x = 0$) and dorsal ($x = 1$) midlines for one of the highly fit parameter sets.

#### 1.2 Gap gene circuit model

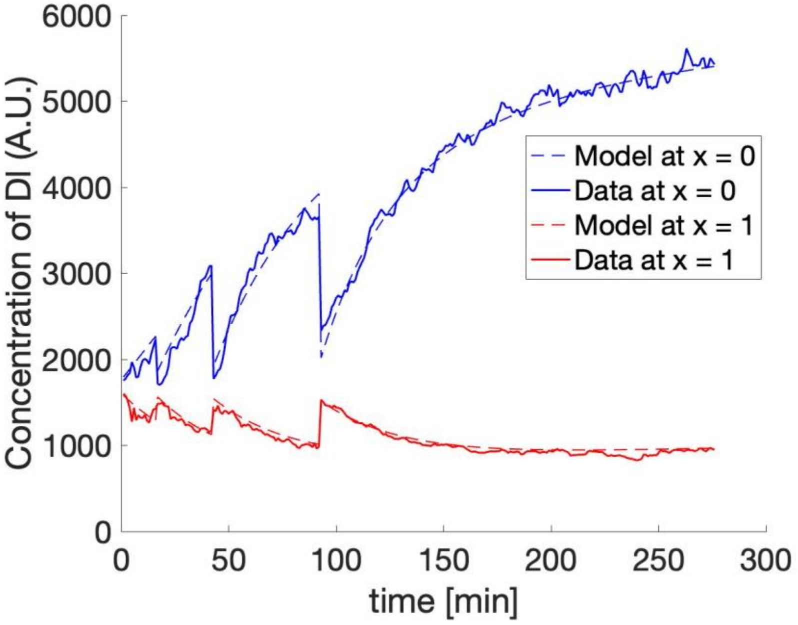


*Figure 1. Model and experimental data plotted for the Dl/Cact model. The plot represents Dl concentration profile from nuclear cycle 11 to 14 at the dorsal and ventral midlines of the early Drosophila embryo.*

##### 1.2.1 Description

In *Drosophila* development, the positions of the 14 body segments in the body of an adult fly are determined to a high degree of precision within ~3hr of development after egg lay (Jäckle et al., 1992). This is accomplished by the Bicoid (Bcd) morphogen that is localized near the anterior pole of the embryo and during early development, it is expressed as a gradient along the anterior-posterior (A-P) direction (Driever and Nusslein-Volhard, 1988). Bcd initiates an asymmetric gene expression cascade in the A-P direction by first activating gap genes that are expressed in broad overlapping domains, followed by pair-rule genes that are expressed in the seven stripes that determine the location of the 14 body segments In this model, the regulatory interactions between the gene regulatory network (GRN) components of the genes involved up to the gap gene part of the cascade are modeled (Manu et al., 2009, Jaeger et al., 2004).

##### 1.2.2 Mathematical model

Gap gene circuit model denotes the intracellular dynamics of protein concentrations along the 58 nuclei in the 35-92% anterior-posterior position during nuclear cycles 13 and 14A. This model for gap gene circuit includes *bcd*, *cad*, *hb*, *Kr*, *kni*, *gt* and the terminal gap gene *tll*. (Manu et al., 2009)

The mathematical model is expressed as follows,

$$\frac{dv_i^a}{dt} = R^a g\left(\sum_{b=1}^{N} T^{ab} v_i^a + m^a v_i^{Bcd} + \sum_{\beta=1}^{N_e} E^{a\beta} v_i^{\beta}(t) + h^a\right) + D^a[(v_{i-1}^a - v_i^a) + (v_{i+1}^a - v_i^a)] - \lambda^a v_i^a \quad (1)$$

where superscripts $a, b \in (1, \ldots, N)$; N is the number of genes, namely *hb*, *Kr*, *kni*, *gt*; superscripts $\beta \in (1, \ldots, N_e)$; $N_e$ is the number of time-varying inputs (Cad and Tll); subscript $i \in (1, \ldots, M)$; and M is the number of nuclei. The three terms on the right side of the equation represent protein synthesis, Fickian diffusion and first order protein degradation, here

$$\mathrm{g(u)} = \frac{1}{2}\left[\left(\frac{\mathrm{u}}{\sqrt{\mathrm{u}^2 + 1}}\right) + 1\right]$$

$g$ is the sigmoidal regulation function, $v_i^a(t)$ is the concentration of the $a^{th}$ species in the $i^{th}$ nucleus at time t, $R^a$ is the maximum synthesis rate of species $a$, $T^{ab}$ and $E^{\alpha\beta}$ are coefficients that weight the transcriptional effect of species b or species β on species a (positive ⇒ activation, negative ⇒ repression), and $h_a$ represents the effect of ubiquitous transcription factors and sets threshold for activation.

##### 1.2.3 Simulation Conditions

The four gap genes; *hb*, *Kr*, *kni*, *gt*; are expressed along the A-P axis, there are 57 nuclei which we discretized space in to 57 1D points, resulting in 232 coupled ordinary differential equations (ODEs). The inputs to the model are: For nuclear cycle 13, ode15s integrates over the ODEs using the initial conditions of *hb*, *Kr*, *kni*, *gt* expression and for nuclear cycle 14A, ode15s uses the simulated results from nuclear cycle 13 as initial conditions. The initial condition for *hb* is experimental data, whereas *Kr, kni*, *gt* are zeros.

##### 1.2.4 Model Results

In Fig. 2, we plot the expression profiles of the four gap genes, namely, *hb*, *Kr, gt*, and *kni,* against model predictions from one of the highly fit (low error) parameter set.

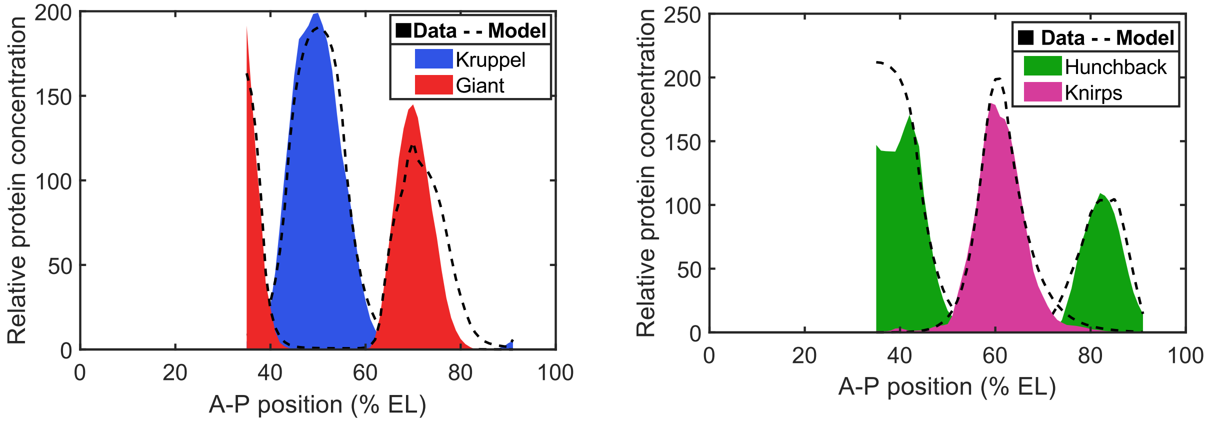


*Figure 2. Concentration profiles of the four gap genes. The plots represent the concentration profiles of Kr, Gt, Hb, and Kni along the anterior-posterior axis based on the conditions specified by the gap gene circuit model.*

#### 1.3 Smad signaling model

##### 1.3.1 Description

Transforming growth factor beta (TGF-β), the founding member of the TGF-β super-family of transforming growth factors, induces Smad signaling, which controls a developmental processes such as embryogenesis, proliferation and apoptosis throughout the body (Aashaq et al., 2022, Kitisin et al., 2007, Huang and Huang, 2005, Schuster and Krieglstein, 2002). TGF-β triggers the signaling cascade by binding to and activating the receptor. Activated receptors phosphorylate Smad2, the receptor-regulated Smad. The phosphorylated Smad2 (pSmad2) forms heteromeric complexes with Smad4, the common Smad. These complexes translocate to the nucleus and regulate gene expression. All the Smad species move dynamically between the cytoplasm and the nucleus, so in presence of TGF-β signaling, Smad2, phospho-Smad2, Smad2-Smad2, and Smad2-Smad4 are present in the nucleus (Massagué, 2012, Schmierer and Hill, 2007).

Schmierer et al. 2008, visualized Smad nucleo-cytoplasmic dynamics with EGFP-tagged-Smad2 through live imaging and quantified the accumulation of Smad species in the nucleus. After 45 mins they added an inhibitor which deactivated the receptors and observed that the nuclear Smad2 concentration decreased as in the absence of TGF-β signaling, Smad2 and Smad4 are inactive and do not form heteromeric complexes, so the only Smad-containing species in the nucleus are Smad2 and Smad4. They used this dynamic nuclear-Smad2 concentration to optimize their mathematical model and found that the Smad2-Smad2 and Smad2-Smad4 complexes have a higher import rate (Complex Import Factor (CIF) > 1) hence they accumulate in the nucleus (Schmierer et al., 2008).

##### 1.3.2 Mathematical model

We used the Retention/Enhanced Complex Import (RECI) model reported by Schmierer et al. (2008), that has 25 state variables and 10 variable parameters. The authors estimated the values of four parameters and optimized the other six. In our simulations, we optimized all 10 parameters.

$$\frac{d[R]}{dt} = -k_{TGF-\beta}[R][TGF-\beta] \qquad (1.)$$

$$\frac{d[TGF-\beta]}{dt} = -k_{TGF-\beta}[R][TGF-\beta] \qquad (2.)$$

$$\frac{d[R^{act}]}{dt} = k_{TGF-\beta}[R][TGF-\beta] - k_{onSB}[R^{act}][SB] + k_{offSB}\left[R^{inact}\right] \qquad (3.)$$

$$\frac{d\left[R^{inact}\right]}{dt} = k_{onSB}[R^{act}][SB] - k_{offSB}\left[R^{inact}\right] \qquad (4.)$$

$$\frac{d[SB]}{dt} = k_{offSB}\left[R^{inact}\right] - k_{onSB}[R^{act}][SB] \qquad (5.)$$

$$\frac{d[S2]_c}{dt} = k_{ex}[S2]_n - k_{in}[S2]_c - k_{phos}[S2]_c[R^{act}] \qquad (6.)$$

$$\frac{d[G]_c}{dt} = k_{ex}[G]_n - k_{in}[G]_c - k_{phos}[G]_c[R^{act}] \qquad (7.)$$

$$\begin{aligned}\frac{d[pS2]_c}{dt} = {} & k_{ex}[pS2]_n - k_{in}[pS2]_c + k_{phos}[S2]_c[R^{act}] \\ & - k_{on}[pS2]_c([S4]_c + 2[pS2]_c + [pG]_c) \\ & + k_{off}([S24]_c + 2[S22]_c + [G2]_c)\end{aligned} \qquad (8.)$$

$$\begin{aligned}\frac{d[pG]_c}{dt} = {} & k_{ex}[pG]_n - k_{in}[pG]_c + k_{phos}[G]_c[R^{act}] \\ & - k_{on}[pG]_c([S4]_c + 2[pS2]_c + [pG]_c) \\ & + k_{off}([G4]_c + 2[G2]_c + [GG]_c)\end{aligned} \qquad (9.)$$

$$\frac{d[S4]_c}{dt} = k_{in}[S4]_n - k_{in}[S4]_c - k_{on}[S4]_c([pS2]_c + [pG]_c) + k_{off}([S24]_c + [G4]_c) \quad (10.)$$

$$\frac{d[S24]_c}{dt} = k_{on}[pS2]_c[S4]_c - k_{off}[S24]_c - k_{in}CIF[S24]_c \quad (11.)$$

$$\frac{d[G4]_c}{dt} = k_{on}[pG]_c[S4]_c - k_{off}[G4]_c - k_{in}CIF[G4]_c \quad (12.)$$

$$\frac{d[S22]_c}{dt} = k_{on}[pS2]_c^2 - k_{off}[S22]_c - k_{in}CIF[S22]_c \quad (13.)$$

$$\frac{d[G2]_c}{dt} = k_{on}[pG]_c[pS2]_c - k_{off}[G2]_c - k_{in}CIF[G2]_c \quad (14.)$$

$$\frac{d[GG]_c}{dt} = k_{on}[pG]_c^2 - k_{off}[GG]_c - k_{in}CIF[GG]_c \quad (15.)$$

$$\frac{d[S2]_n}{dt} = k_{in}[S2]_c - k_{ex}[S2]_n + k_{dephos}[pS2]_n[PPase] \quad (16.)$$

$$\frac{d[G]_n}{dt} = k_{in}[G]_c - k_{ex}[G]_n + k_{dephos}[pG]_n[PPase] \quad (17.)$$

$$\frac{d[pS2]_n}{dt} = k_{in}[pS2]_c - k_{ex}[pS2]_n - k_{dephos}[pS2]_n[PPase] - k_{on}[pS2]_n([S4]_n + 2[pS2]_n + [pG]_n) + k_{off}([S24]_c + 2[S22]_c + [G2]_c) \quad (18.)$$

$$\frac{d[pG]_n}{dt} = k_{in}[pG]_c - k_{ex}[pG]_n - k_{dephos}[pG]_n[PPase] - k_{on}[pG]_n([S4]_n + [pS2]_n + 2[pG]_n) + k_{off}([G4]_c + 2[G2]_c + [GG]_c) \quad (19.)$$

$$\frac{d[S4]_n}{dt} = k_{in}[S4]_c - k_{in}[S4]_n - k_{on}[S4]_n([pS2]_n + [pG]_n) + k_{off}([S24]_n + [G4]_n) \quad (20.)$$

$$\frac{d[S24]_n}{dt} = k_{on}[pS2]_n[S4]_n - k_{off}[S24]_n + k_{in}CIF[S24]_c \quad (21.)$$

$$\frac{d[G4]_n}{dt} = k_{on}[pG]_n[S4]_n - k_{off}[G4]_n + k_{in}CIF[G4]_c \quad (22.)$$

$$\frac{d[S22]_n}{dt} = k_{on}[pS2]_n^2 - k_{off}[S22]_n + k_{in}CIF[S22]_c \quad (23.)$$

$$\frac{d[G2]_n}{dt} = k_{on}[pG]_n[pS2]_n - k_{off}[G2]_n + k_{in}CIF[G2]_c \quad (24.)$$

$$\frac{d[GG]_n}{dt} = k_{on}[pG]_n^2 - k_{off}[GG]_n + k_{in}CIF[GG]_c \quad (25.)$$

where, subscripts $n$ and $c$ represent nucleus and cytoplasm, R represents the unbound receptor, superscripts $act$ and $inact$ represent TGF-β activated receptor and SB (inhibitor) bound receptor, SB is the inhibitor, S2 is Smad2, pS2 is Phospho-Smad2, S4 is Smad4, S24 is Smad2/Smad4 complex, S22 is Samd2/Smad2 complex, PPase is Phosphatase, G is EGFP-Smad2, pG is the phosphorylated EGFP-Smad2, G4 is the EGFP-Smad2/Smad4 complex, G2 is EGFP-Smad2/Smad2 complex, and GG is the EGFP-Smad2/EGFP-Smad2 complex.

The parameter $k$ stands for rate constants in the mathematical model. The subscripts denote the process that the two reacting species are involved in. $k_{\mathrm{TGF-\beta}}$ denotes the activation of receptors in the presence of TGF-β. The parameters $k_{on}$ and $k_{off}$ denote binding and unbinding rates for Phospho-Smad2 and Smad4, while $k_{onSB}$ and $k_{offSB}$ denote binding and unbinding rates for activated receptor ($R^{act}$) and the inhibitor (SB). The parameters $k_{in}$ and $k_{out}$ denote the rate at which complexes translocate in and out of the nucleus and, $k_{dephos}$ denotes the dephosphorylation rate of Phospho-Smad2 by Phosphatase. CIF is the complex import factor, which is the fold difference between the import rate of Smad complexes and import rate of monomeric Smads.

##### 1.3.3 Simulation conditions

We used figure 2D from Schmierer et al. (2008), to obtain the nuclear EGFP-Smad2 concentration with time. The model has two parts: (1) at time t = 0, $T$GF-β is added to the system, so that signaling is on, and (2) at time t = 45 min, inhibitor is added to the system, which turns off TGF-β signaling. For part (1), we simulated the model equations for 45 mins with no inhibitor. For part (2), we added inhibitor to the system and simulated with concentration at $t$ = 45 min as initial condition. We simulated this system of equations using ode15s in MATLAB with randomly generated parameter sets and calculated the sum of squared errors between the modeled profile and nuclear EGFP-Smad2 concentrations reported in Figure 2D from Schmierer et al. (2008) and normalized it with the error bars reported in Figure 2D from Schmierer et al. (2008). The initial concentrations of the species reported are as follows:

| Species | Nucleus (nM) | Cytoplasm (nM) |
|---|---|---|
| Smad 2/3 + EGFP-Smad2 | 28.5 + 28.5 | 60.6 + 60.6 |
| Smad4 | 50.8 | 50.8 |
| Other Smads | 0 | 0 |
| Inactive receptors | 1 | |
| Phosphatase | | 1 |
| $T$GF-β | | 0.066 |
| SB (inhibitor) | | 10,000 |

The rest of the Smad species have a zero initial value.

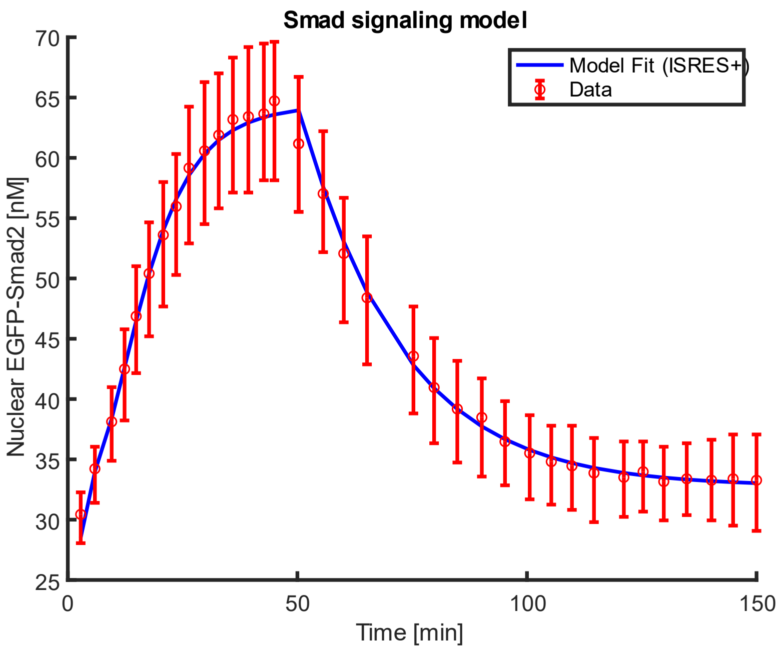


*Figure 3. EGFP-Smad2 concentration profile*

##### 1.3.4 Model results

In the Fig. 3, we plot the concentration of the nuclear EGFP-Smad2 complex data against the predicted value by one a high fit model parameter set. Please refer to Fig 2D in the Schmierer et al. (2008) for comparison.

### 2. Derivation of Linstep and Newton step

#### 2.1 Linstep

The Linstep fitting equation may be represented as follows,

$$\boldsymbol{f_i} = a_0 + a_1\boldsymbol{\theta_1^i} + a_2\boldsymbol{\theta_2^i} + \ldots + a_n\boldsymbol{\theta_n^i}$$

$$\boldsymbol{f_i} = a_0 + \sum_{j=1}^{n} a_j\theta_j^i$$

where, $i = 1 \ldots N$ are the number of individuals used in the fitting procedure.

$$\begin{bmatrix} f_1 \\ f_2 \\ \vdots \\ f_N \end{bmatrix} = \begin{bmatrix} 1 & \theta_1^1 & \theta_2^1 & \cdots & \theta_n^1 \\ 1 & \theta_1^2 & \theta_1^1 & \cdots & \theta_n^2 \\ 1 & \theta_1^3 & \theta_1^1 & \cdots & \theta_n^3 \\ \vdots & \vdots & \vdots & \vdots & \vdots \\ 1 & \theta_1^N & \theta_1^N & \cdots & \theta_n^N \end{bmatrix} \begin{bmatrix} a_0 \\ a_1 \\ \vdots \\ a_N \end{bmatrix}$$

$$a = (M^T M^{-1}) M^T F$$

#### 2.2 Newton step

The Newton step fitting equation may be represented as follows,

$$\boldsymbol{f}(\boldsymbol{\theta}^i) = \boldsymbol{f}(\boldsymbol{\theta}^\circ) + \boldsymbol{\nabla f}_{\boldsymbol{\theta}^\circ}\,\Delta\boldsymbol{\theta} + \frac{1}{2}\Delta\boldsymbol{\theta}^{\mathrm{T}}\mathbf{H_0}\Delta\boldsymbol{\theta}$$

where $\Delta\boldsymbol{\theta} = \boldsymbol{\theta}^{i} - \boldsymbol{\theta}^{\circ}$, where $\boldsymbol{\theta}^{\circ}$ and $\boldsymbol{\theta}^{\boldsymbol{i}}$ refers to the fittest individual in the population and to individual $i$, and $\mathbf{H_0}$ is the Hessian at the $\boldsymbol{\theta} = \boldsymbol{\theta}^{\circ}$.

The third term can be written as,

$$\frac{1}{2}\Delta\boldsymbol{\theta}^{\mathrm{T}}\mathbf{H_0}\Delta\boldsymbol{\theta} = [\Delta\theta_1 \quad \Delta\theta_2 \quad \cdots \quad \Delta\theta_n]\begin{bmatrix} H_{11} & H_{12} & H_{13} & \cdots & H_{1n} \\ H_{21} & H_{22} & H_{23} & \cdots & H_{2n} \\ H_{31} & H_{32} & H_{33} & \cdots & H_{3n} \\ \vdots & \vdots & \vdots & \vdots & \vdots \\ H_{n1} & H_{n2} & H_{n3} & \cdots & H_{nn} \end{bmatrix}\begin{bmatrix} \Delta\theta_1 \\ \Delta\theta_2 \\ \vdots \\ \Delta\theta_n \end{bmatrix}$$

$$= \frac{1}{2}\sum_{i=1}^{n}\sum_{j=1}^{n} H_{jk}\Delta\theta_j\Delta\theta_k$$

where $H_{jk} = \frac{\partial^2 f}{\partial\theta_j\partial\theta_k}$ and $\Delta\theta_k = \theta_k^i - \theta_k^{\circ}$. The term can be simplified as follows,

$$\frac{1}{2}\sum_{i=1}^{n}\sum_{j=1}^{n} H_{jk}\Delta\theta_j\Delta\theta_k = \frac{1}{2}\sum_{i=1}^{n}\sum_{j=1}^{n} H_{jk}\left(\theta_j^i - \theta_j^{\circ}\right)\left(\theta_k^i - \theta_k^{\circ}\right)$$

$$= \frac{1}{2}\sum_{i=1}^{n}\sum_{j=1}^{n} H_{jk}\left(\theta_j^i\theta_k^i - \theta_j^i\theta_k^{\circ} - \theta_j^{\circ}\theta_k^i + \theta_j^{\circ}\theta_k^{\circ}\right)$$

$$= \frac{1}{2}\sum_{i=1}^{n}\sum_{j=1}^{n}\left(H_{jk}\theta_j^i\theta_k^i + a_{jk}\theta_j^i + b_{jk}\theta_j^{\circ}\theta_k^i + c_{jk}\right)$$

where $a_{jk} = -H_{jk}\theta_k^{\circ}$, $b_{jk} = -H_{jk}\theta_j^{\circ}$, and $c_{jk} = -H_{jk}\theta_j^{\circ}\theta_k^{\circ}$.

The second term may be written as,

$$\boldsymbol{\nabla f}\,\Delta\boldsymbol{\theta} = \left[f_{\theta_1} \quad f_{\theta_2} \quad \cdots \quad f_{\theta_n}\right]\begin{bmatrix} (\theta_1^i - \theta_1^{\circ}) \\ (\theta_2^i - \theta_2^{\circ}) \\ \vdots \\ (\theta_n^i - \theta_n^{\circ}) \end{bmatrix}$$

$$= \sum_{j=1}^{n} f_{\theta_j}(\theta_j^i - \theta_j^\circ)$$

Both the terms can be put together as follows,

$$\boldsymbol{f_i} = \frac{1}{2}\sum_{j=1}^{n}\sum_{k=1}^{n} H_{jk}\theta_j^i\theta_k^i + \sum_{j=1}^{n} g_j\theta_j^i + c$$

where $\boldsymbol{g_j} = f_{\theta_j} - \sum_{k=1}^{n} H_{jk}\theta_k^\circ$ and $c = \sum_{j=1}^{n} f_{\theta_j^\circ} - \sum_{j=1}^{n} f_{\theta_j}\theta_j^\circ + \frac{1}{2}\sum_{j=1}^{n}\sum_{k=1}^{n} H_{jk}\theta_j^\circ\theta_k^\circ$. Thus, the elements of the Hessian matrix and the elements of the gradient can be estimated by inverting the matrix consisting of combinations of $\boldsymbol{\theta^i}$ on the right hand side similar to Linstep.

#### 2.3 A full Newton step leads all individuals considered in the Hessian calculation to move to the same point

Differentiating the second order fit equation,

$$\boldsymbol{f}(\boldsymbol{\theta}^i) = \boldsymbol{f}(\boldsymbol{\theta}^\circ) + \boldsymbol{\nabla f_{\theta^\circ}}\,\Delta\boldsymbol{\theta} + \frac{1}{2}\Delta\boldsymbol{\theta}^{\mathrm{T}}\mathbf{H_0}\Delta\boldsymbol{\theta}$$

with respect to $\theta$ gives,

$$0 = (\boldsymbol{\nabla f}^\circ)^T + \mathbf{H}^\circ\Delta\theta$$

Therefore,

$$\boldsymbol{\theta^i} = \boldsymbol{\theta}^\circ - (\mathbf{H}^\circ)^{-1}[(\boldsymbol{\nabla f}^\circ)^T]$$

Using the definition of $g_j$ from the above derivation, and rearranging give,

$$f_{\theta_j} = g_j + \sum_{k=1}^{n} H_{jk}\theta_k^\circ$$

we can rewrite the equation as,

$$\boldsymbol{\theta}^{\boldsymbol{i}} = \boldsymbol{\theta}^{\circ} - (\mathbf{H}^{\circ})^{-1}[\boldsymbol{g} + \mathbf{H}^{\circ}\theta^{\circ}]$$

$$= (\mathbf{H}^{\circ})^{-1}\boldsymbol{g}$$
